# LDpred2: better, faster, stronger

**DOI:** 10.1101/2020.04.28.066720

**Authors:** Florian Privé, Julyan Arbel, Bjarni J. Vilhjálmsson

**Author notes:** To whom correspondence should be addressed. Contacts.

## Abstract

Polygenic scores have become a central tool in human genetics research. LDpred is a popular method for deriving polygenic scores based on summary statistics and a matrix of correlation between genetic variants. However, LDpred has limitations that may reduce its predictive performance. Here we present LDpred2, a new version of LDpred that addresses these issues. We also provide two new options in LDpred2: a “sparse” option that can learn effects that are exactly 0, and an “auto” option that directly learns the two LDpred parameters from data. We benchmark predictive performance of LDpred2 against the previous version on simulated and real data, demonstrating substantial improvements in robustness and predictive accuracy compared to LDpred1. We then show that LDpred2 also outperforms other polygenic score methods recently developed, with a mean AUC over the 8 real traits analyzed here of 65.1%, compared to 63.8% for lassosum, 62.9% for PRS-CS and 61.5% for SBayesR. Note that, in contrast to what was recommended in the first version of this paper, we now recommend to run LDpred2 genome-wide instead of per chromosome. LDpred2 is implemented in R package bigsnpr.

## 1 Introduction

In recent years the use of polygenic scores (PGS) has become widespread. A PGS aggregates (risk) effects across many genetic variants into a single predictive score. These scores have proven useful for studying the genetic architecture and relationships between diseases and traits (Purcell *et al*. 2009; Kong *et al*. 2018). Moreover, there are high hopes for using these scores in clinical practice to improve disease risk estimates and predictive accuracy. The heritability, i.e. the proportion of phenotypic variance that is attributable to genetics, determines an upper limit on the predictive performance of PGS and thus their value as a predictive tool. Nevertheless, a number of studies have explored the use of PGS in clinical settings (Pashayan *et al*. 2015; Willoughby *et al*. 2019; Abraham *et al*. 2019). PGS are also extensively used in epidemiology and economics as predictive variables of interest (Musliner *et al*. 2015; Horsdal *et al*. 2019; Barth *et al*. 2020; Harden and Koellinger 2020). For example, a recently derived PGS for education attainment has been one of the most predictive variables in behavioural sciences so far (Allegrini *et al*. 2019).

LDpred is a popular method for deriving polygenic scores based on summary statistics and a Linkage Disequilibrium (LD) matrix only (Vilhjálmsson *et al*. 2015). However, LDpred has several limitations that may reduce its predictive performance. The non-infinitesimal version of LDpred, which assumes there is a proportion *p* of variants that are causal, is a Gibbs sampler and is particularly sensitive to model misspecification when applied to summary statistics with large sample sizes. It is also unstable in long-range LD regions such as the human leukocyte antigen (HLA) region of chromosome 6. This issue has led to the removal of such regions from analyses (Marquez-Luna *et al*. 2020; Lloyd-Jones *et al*. 2019), which is unfortunate since this region of the genome contains many known disease-associated variants, particularly with autoimmune diseases and psychiatric disorders (Mokhtari and Lachman 2016; Matzaraki *et al*. 2017).

Here, we present LDpred2, a new version of LDpred that addresses these issues while markedly improving its computational efficiency, allowing exploring a larger grid of parameters in the same computational time as LDpred1. We provide this faster and more robust implementation of LDpred in R package bigsnpr (Privé *et al*. 2018). We also provide two new options in LDpred2. First, we provide a “sparse” option, where LDpred2 truly fits some effects to zero, therefore providing a sparse vector of effects. Second, we also provide an “auto” option, where LDpred2 automatically estimates the sparsity *p* and the SNP heritability *h*^2^, and therefore does not require validation data to tune hyper-parameters. We show that LDpred2 provides higher predictive performance than LDpred1 (LDpred v1.0.0), especially when there are causal variants in long-range LD regions, when the proportion of causal variants is small, and when GWAS sample size is large. We also show that the new sparse option performs equally well as the non-sparse version, enabling LDpred2 to provide sparse effects without losing prediction accuracy. Moreover, LDpred2-auto, which does not require any validation set, performs almost as well as the main LDpred2 model that tunes hyper-parameters from a grid in a validation set, provided some quality control is performed on the summary statistics.

## 2 Results

### Overview of methods

Here we present LDpred2, a new version of LDpred (Vilhjálmsson *et al*. 2015). LDpred2 has 3 options: 1) LDpred2-inf, which provides an analytical solution under the infinitesimal model of LDpred1; 2) LDpred2-grid (or simply LDpred2) that is the main LDpred model, where a grid of values for hyper-parameters *p* (the proportion of causal variants), *h*^2^ (the SNP heritability), and possibly the sparsity option (as a third hyper-parameter) are tuned using a validation set; 3) LDpred2-auto, which automatically estimates *p* and *h*^2^ from data and therefore is free of hyper-parameters to tune. Note that the sparse option in LDpred2-grid slightly modifies the Gibbs sampler in LDpred2 to be able to fit effects that are exactly 0. For users who do not particularly care about sparsity, they can still tune this parameter in LDpred2-grid to explore a larger parameter space. For this paper, in order to show the effect of this sparse option on predictive performance, we deliberately separate the LDpred2-grid method in two “different” methods that we call LDpred2-grid-nosp (sparse option always disabled) and LDpred2-grid-sp (sparse option always enabled) in the results. As a recall, LDpred v1 has two options: LDpred1-inf, and LDpred1-grid where only *p* is tuned while *h*^2^ is estimated from constrained LD score regression (Bulik-Sullivan *et al*. 2015).

We make use of the UK Biobank data to compare the two versions of LDpred using several simulation scenarios to understand the expected impact of using the new version of LDpred. We also compare these two versions of LDpred using eight case-control phenotypes of major interest and for which there are published external summary statistics available and a substantial number of cases in the UK Biobank data. We also compare running LDpred2 per chromosome (“perchr”, including choosing optimal hyper-paramaters) or genome-wide (“gwide”). Finally, we compare LDpred2 to several other methods: Clumping and Thresholding (C+T), Stacked C+T (SCT), lassosum, PRS-CS and SBayesR (Privé *et al*. 2019; Mak *et al*. 2017; Ge *et al*. 2019; Lloyd-Jones *et al*. 2019). Area Under the ROC Curve (AUC) values are reported.

### Simulations

Figure 1 presents the simulation results comparing LDpred1 (v1.0.0 as implemented by Vilhjálmsson *et al*. (2015)) with the new LDpred2 (as implemented in R package bigsnpr). Seven simulation scenarios are used, each repeated 10 times. In the first four simulation scenarios, a heritability *h*^2^ of 40% and a prevalence of 15% are used. We simulate 300, 3000, 30,000 or 300,000 causal variants anywhere on the genome. In these scenarios, infinitesimal models perform similarly. When testing a grid of hyper-parameters, LDpred2 performs substantially better than LDpred1 in the cases where the number of causal variants is small, i.e. in the case of 300 or 3000 causal variants (*p*=2.7e-4 and 2.7e-3). For example, in simulations with 300 causal variants, a mean AUC of 73.6% is obtained with LDpred1 while a value of 81.7% is obtained with LDpred2. In these scenarios, all non-infinitesimal models of LDpred2 perform equally well. As expected, LDpred1 performs poorly in HLA scenarios, i.e. when causal variants are located in a long-range LD region. In contrast, all LDpred2 models perform well in these two HLA scenarios, but LDpred2-auto performs slightly worse than the other LDpred2 models in these extreme scenarios.

**Figure 1:**
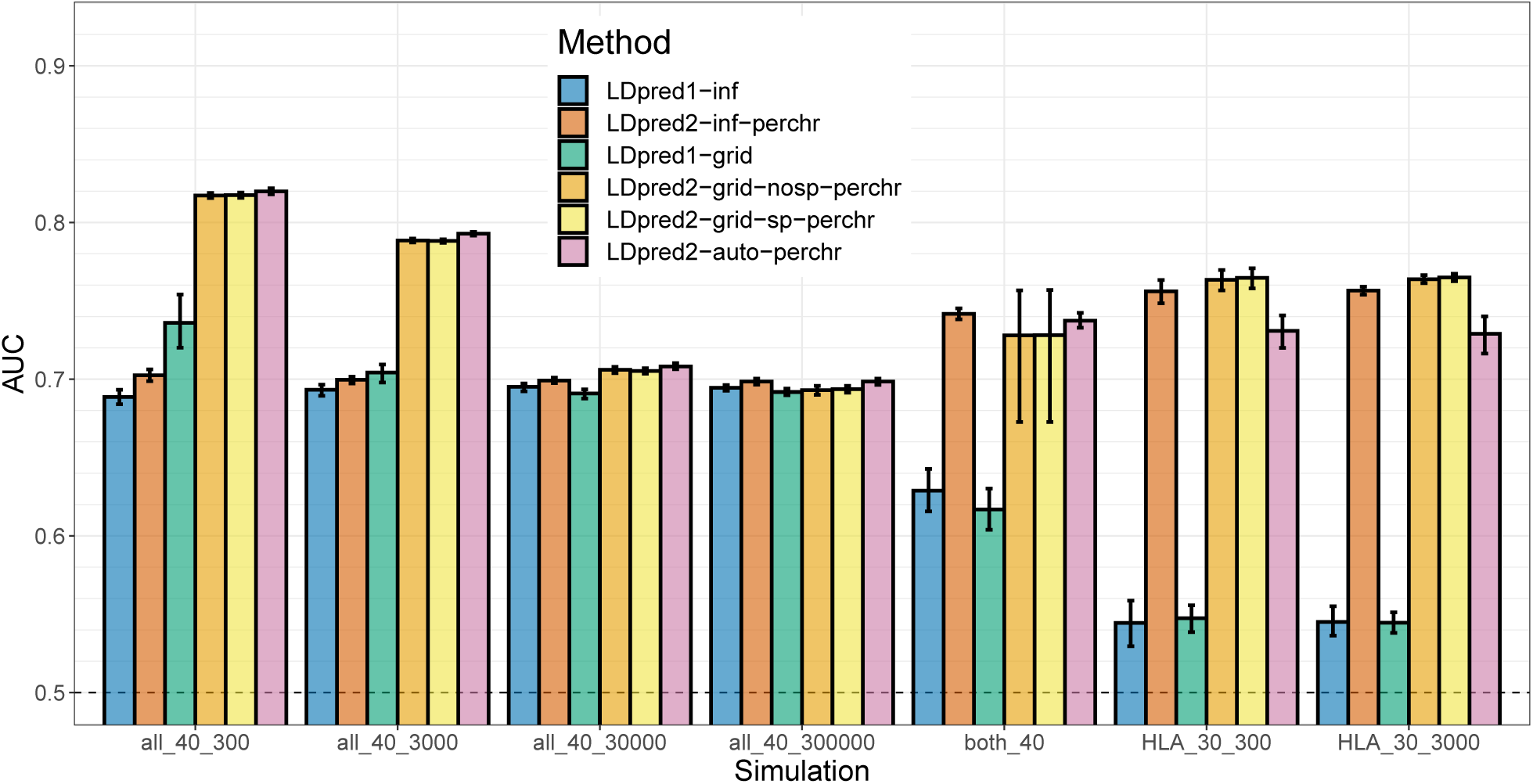
Two variants of LDpred1 are compared with four variants of LDpred2 (run per chromosome) in the seven simulation scenarios summarized in table 1. Briefly, the first part of the scenario name corresponds to the location of causal variants, the second part is the heritability (in %), the third part is the number of causal variants, and the prevalence is always 15%. Bars present the mean and 95% CI of 10,000 non-parametric bootstrap replicates of the mean AUC of 10 simulations for each scenario. Corresponding values are reported in table S1.

Figure 2 presents the simulation results comparing LDpred1-grid and LDpred2-grid when varying the GWAS sample size from 10,000 to 300,000 in the simulation scenario with 3000 causal variants anywhere on the genome (“all_40_3000”). LDpred1-grid performs equally well as LDpred2-grid (when LDpred2 is run genome-wide) for the smallest GWAS sample sizes. Yet, LDpred1-grid starts providing lower predictive performance than LDpred2-grid when GWAS sample size becomes larger. For example, in the scenario with a sample size of 120,000, LDpred1 provides a mean AUC of 70.7% while LDpred2 provides a mean AUC of 74.6%. Moreover, LDpred1 even performs slightly worse with a mean AUC of 70.4% when the sample size is increased to 300,000. In contrast, the performance of LDpred2 continues to increase steadily to 79.1% towards the maximum achievable AUC of 82.5% for this scenario with a heritability of 40% and a prevalence of 15% (Wray *et al*. 2010).

**Table 1:** The seven simulations scenarios. N: GWAS sample size.

| Simulation | Number and location of causal variants | Heritability | N (x1000) | Prevalence |
| --- | --- | --- | --- | --- |
| all_40_300 | 300 anywhere on the genome | 40% | 300 | 15% |
| all_40_3000 | 3000 anywhere on the genome | 40% | 10, 20, 50, 120, 300 | 15% |
| all_40_30000 | 30,000 anywhere on the genome | 40% | 300 | 15% |
| all_40_300000 | 300,000 anywhere on the genome | 40% | 300 | 15% |
| HLA_30_300 | 300 in the HLA region | 30% | 300 | 15% |
| HLA_30_3000 | 3000 in the HLA region | 30% | 300 | 15% |
| both_40 | 10,000 anywhere on the genome + 300 in the HLA region | 20% + 20% | 300 | 15% |

**Figure 2:**
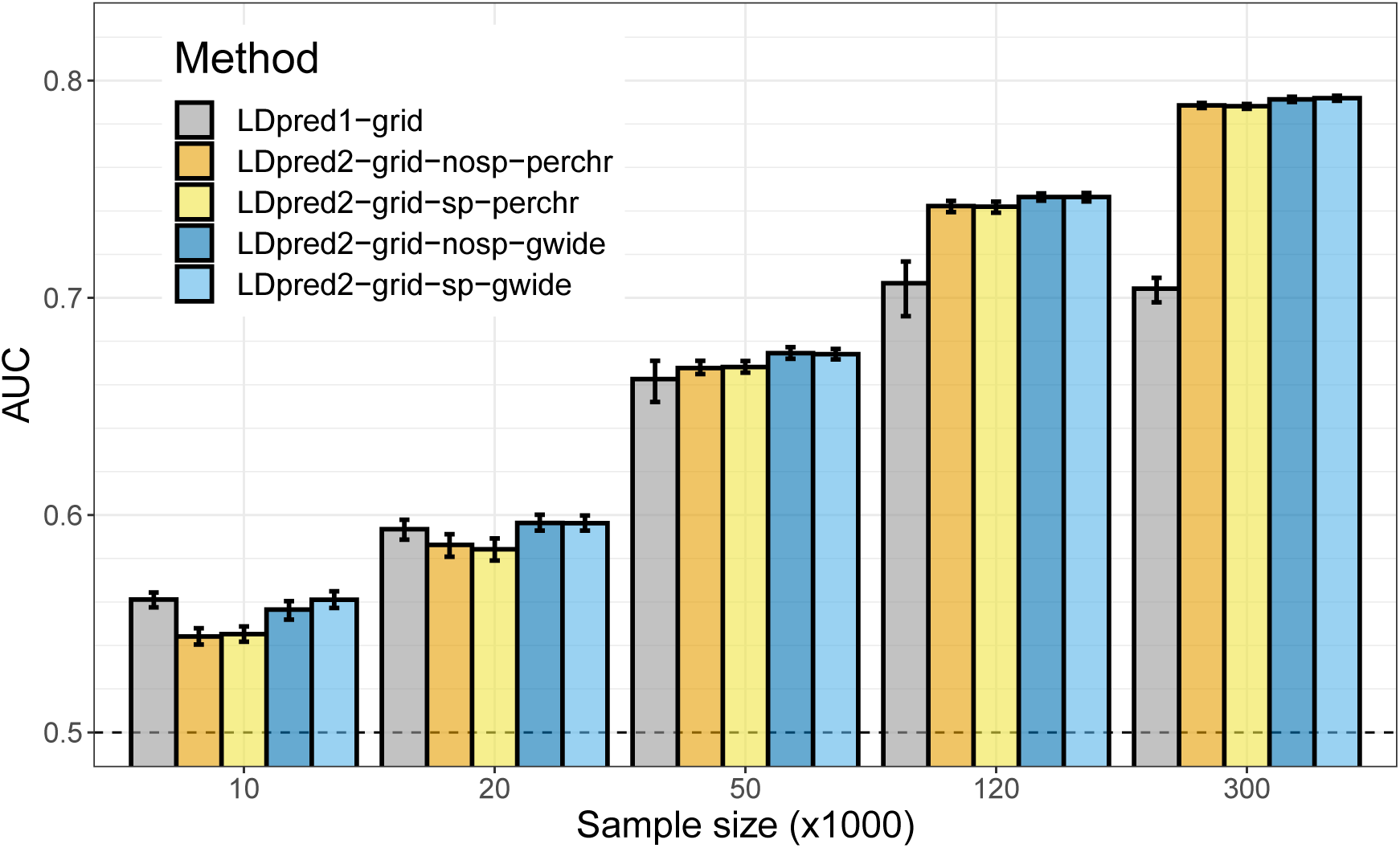
LDpred1-grid is compared to LDpred2-grid when varying GWAS sample size in scenario “all_40_3000”. Bars present the mean and 95% CI of 10,000 non-parametric bootstrap replicates of the mean AUC of 10 simulations for each scenario. Corresponding values are reported in table S2.

### Real data

Figure 3 presents the results of real data applications comparing LDpred1 (v1.0.0 as implemented by Vilhjálmsson *et al*. (2015)) with the new LDpred2 (as implemented in R package bigsnpr) when run genome-wide. Eight case-control phenotypes are used, summarized in table 2. For BRCA, CAD, MDD, PRCA, T1D and T2D, LDpred2-inf and LDpred2-grid perform much better than LDpred1-inf and LDpred1-grid respectively. For example, for BRCA, AUC improves from 58.9% with LDpred1-grid to 65.5% with LDpred2-grid, and from 57.4% to 78.4% for T1D. For Asthma and RA, predictive performance of LDpred1 and LDpred2 are similar. As in simulations, the sparse version of LDpred2-grid performs as well as the non-sparse version. Sparsity of resulting effects ranges from 19.3% for RA to 54.4% for Asthma (Table S5). Figure 4 presents the results of real data applications comparing LDpred2 to several other PGS methods. LDpred2 performs best for all phenotypes, lassosum performs relatively well for all phenotypes, PRS-CS always performs slightly worse than lassosum, and SBayesR performs as well as LDpred2 for BRCA, MDD, PRCA and T2D, but severely underperforms for the autoimmune diseases (T1D and RA). For example,

**Table 2:** Summary of external GWAS summary statistics used. The GWAS sample size is the number of cases / controls in the GWAS. The mean chi-squared statistics *χ*^2^ and the number of hits (*p <* 5 ·10^−8^) are reported after restricting to HapMap3 variants and matching with the UKBB data. Independent hits are the hits remaining after clumping at *r*^2^ > 0.01 within 10 Mbp.

| Trait | GWAS citation | GWAS sample size | # GWAS variants | # matched variants | Mean $\chi^2$ | # hits | # indep hits |
| --- | --- | --- | --- | --- | --- | --- | --- |
| Breast cancer (BRCA) | Michailidou *et al.* (2017) | 137,045 / 119,078 | 11,792,542 | 1,114,424 | 1.70 | 2941 | 157 |
| Rheumatoid arthritis (RA) | Okada *et al.* (2014) | 29,880 / 73,758 | 9,739,303 | 656,087 | 2.10 | 3663 | 75 |
| Type 1 diabetes (T1D) | Censin *et al.* (2017) | 5913 / 8828 | 8,996,866 | 514,420 | 1.93 | 1902 | 39 |
| Type 2 diabetes (T2D) | Scott *et al.* (2017) | 26,676 / 132,532 | 12,056,346 | 1,108,760 | 1.24 | 388 | 39 |
| Prostate cancer (PRCA) | Schumacher *et al.* (2018) | 79,148 / 61,106 | 20,370,946 | 1,115,688 | 1.55 | 2762 | 137 |
| Depression (MDD) | Wray *et al.* (2018) | 59,851 / 113,154 | 13,554,550 | 1,103,440 | 1.27 | 166 | 5 |
| Coronary artery disease (CAD) | Nikpay *et al.* (2015) | 60,801 / 123,504 | 9,455,778 | 1,108,313 | 1.13 | 352 | 37 |
| Asthma | Demènais *et al.* (2018) | 19,954 / 107,715 | 2,001,280 | 980,430 | 1.10 | 365 | 15 |

**Figure 3:**
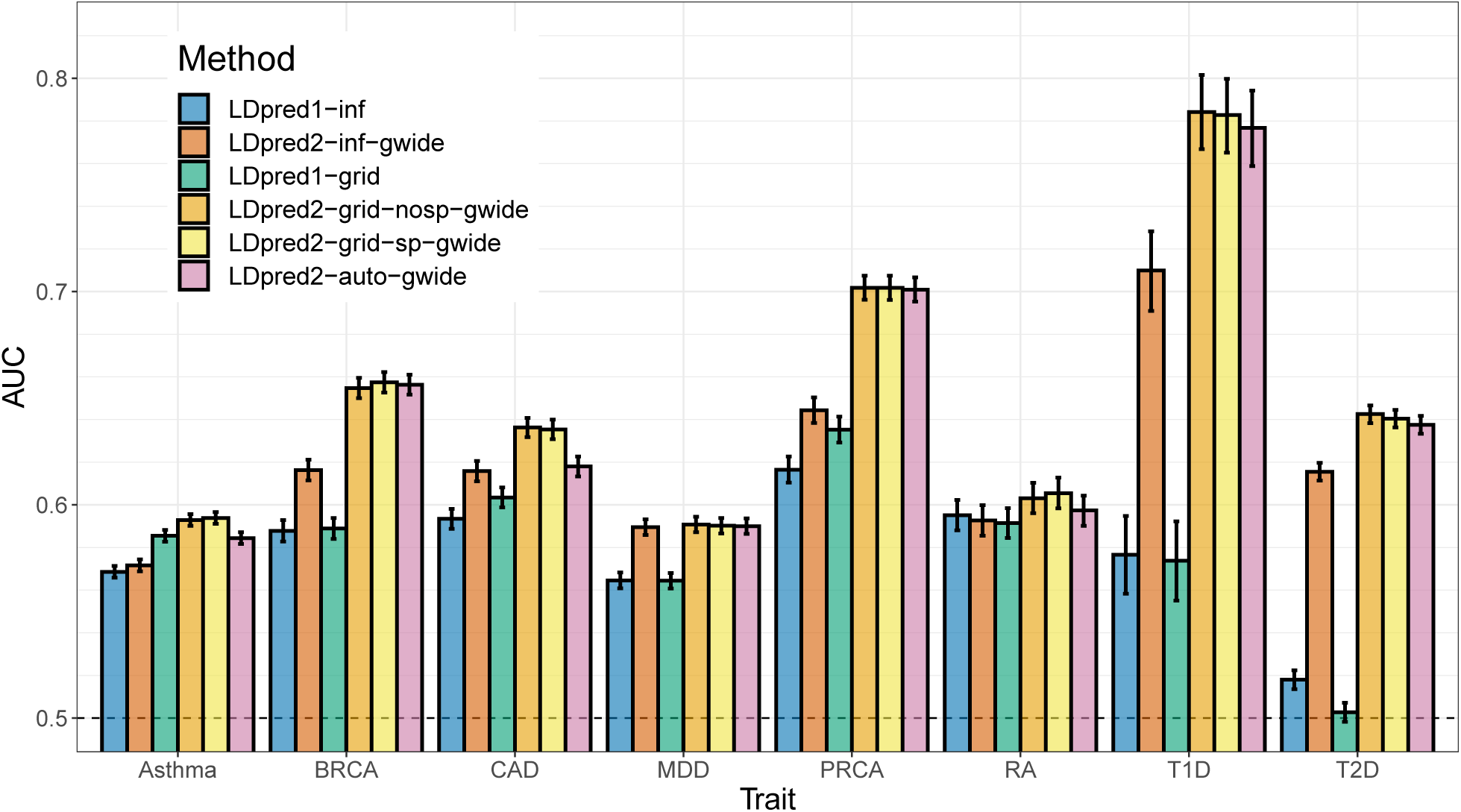
Two variants of LDpred1 are compared with four variants of LDpred2 (run genome-wide) in the real data applications using published external summary statistics. Bars present AUC values on the test set of UKBB (mean and 95% CI from 10,000 bootstrap samples). Corresponding values are reported in table S3.

**Figure 4:**
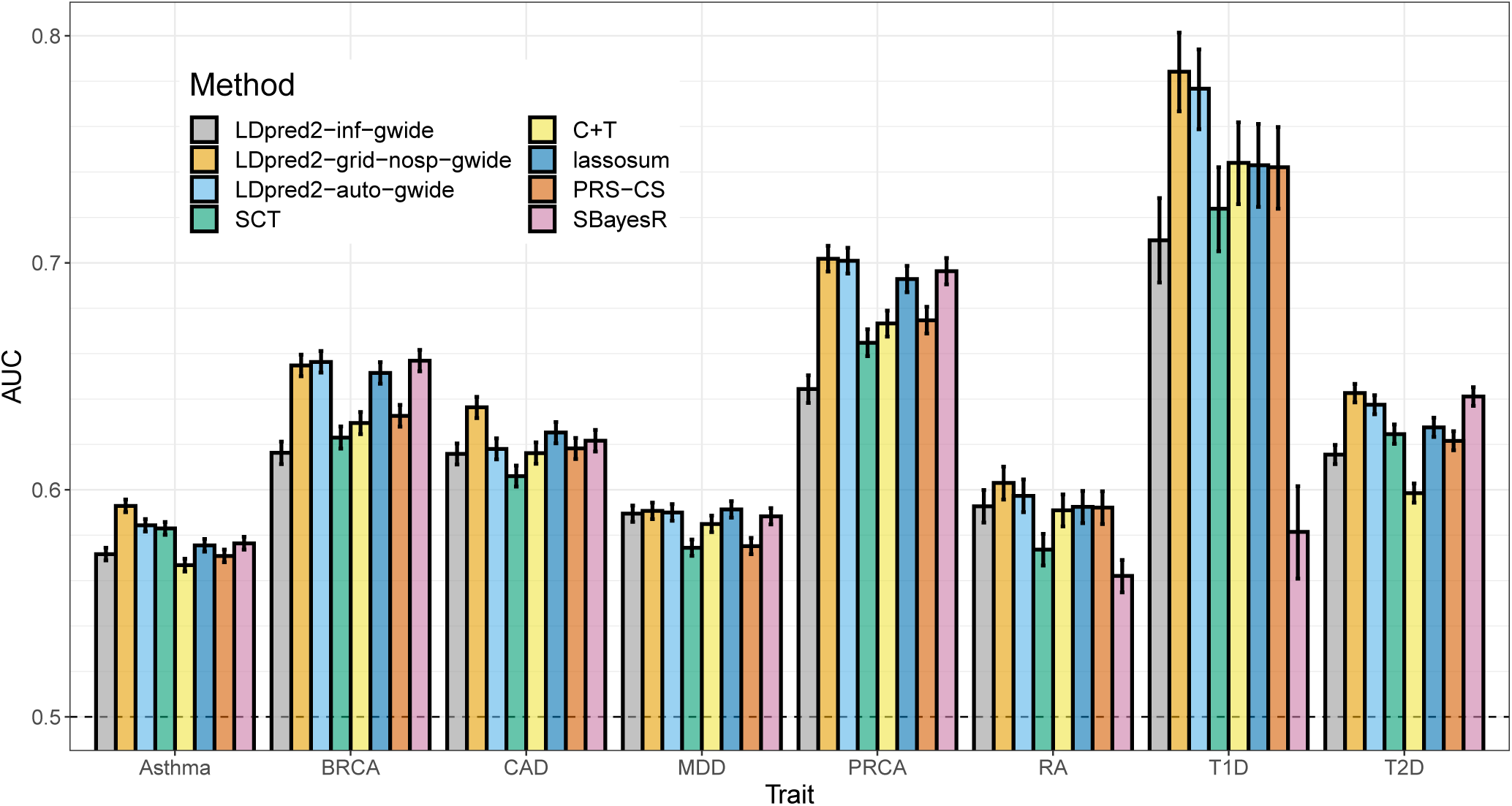
LDpred2 is compared with C+T, SCT, lassosum, PRS-CS and SBayesR in the real data applications using published external summary statistics. Bars present AUC values on the test set of UKBB (mean and 95% CI from 10,000 bootstrap samples). Corresponding values are reported in table S4.

SBayesR provides an AUC of 58.1% for T1D compared to 78.4% with LDpred2-grid.

Figure S1 presents the results of real data applications comparing “grid” models with their “auto” counterparts, i.e. models that directly estimate parameters from the data and do not require tuning hyper-parameters from a grid using a validation set. We remind readers that SBayesR is such an “auto” model, but does not have any “grid” counterpart. All “grid” models usually perform better than their “auto” counterpart. LDpred2-auto performs similarly as LDpred2-grid, except for CAD (AUC of 61.8% vs 63.6%) and to a lesser extent for Asthma (AUC of 58.4% vs 59.3%). Similarly, lassosum-auto performs similarly as lassosum, expect for MDD (AUC of 54.9% vs 59.4%). Similarly, PRS-CS-auto also underperforms for MDD (AUC of 53.9% for PRS-CS-auto vs 57.5% for PRS-CS). In contrast, SBayesR does not underperforms for MDD but severely underperforms for T1D and RA, as discussed in the previous paragraph.

### Running LDpred2 per chromosome or genome-wide

Most results show that it is beneficial to run LDpred2 genome-wide instead of per chromosome, contrary to was recommended in the first version of this paper. Figure S2 shows that it can be beneficial to run all LDpred2 models genome-wide rather than per chromosome, especially when GWAS sample size is small.

For example, in the simulation scenario with the sample size of 20,000, switching running LDpred2 per chromosome to running it genome-wide improves AUC of LDpred2-inf from 57.2% to 57.7%, LDpred2-grid from 58.6% to 59.6%, and LDpred2-auto from 57.3% to 59.3%. Note that the simulation scenario used to produce results in figure S2 is assuming the same genome-wide disease architecture, i.e. 3000 causal variants located anywhere on the genome and whose effects are drawn from the same normal distribution. However, figure S3 also shows that it can be beneficial to run all LDpred2 models genome-wide rather than per chromosome for real traits as well. Indeed, except for LDpred2-inf applied to T1D, all other cases show similar or better performance when running LDpred2 genome-wide. For example, running LDpred2-grid genome-wide rather than per chromosome is particularly beneficial for CAD (AUC of 63.6% vs 61.5%) and PRCA (AUC of 70.2% vs 68.2%).

## 3 Discussion

The previous version of LDpred has been widely used and has the potential to provide polygenic models with good predictive performance (Khera *et al*. 2018). Yet, it has instability issues that have been pointed out (Marquez-Luna *et al*. 2020; Lloyd-Jones *et al*. 2019) and likely contributed to discrepancies in reported predictive accuracy of LDpred (Choi and O’Reilly 2019; Ge *et al*. 2019; Chun *et al*. 2020). We have therefore implemented a new version of LDpred, LDpred2, to address these issues. We show that LDpred2 is more stable and provides higher predictive performance than LDpred1, particularly when handling long-range LD regions, less polygenic traits, and large GWAS sample sizes. We hypothesize that LDpred1 does not use a LD window size that is large enough to account for long-range LD such as in the HLA region. In LDpred2, we use a window size of 3 cM, which is larger than the default value used in LDpred1. This enables LDpred2 to work well even when causal variants are located in long-range LD regions. We argue against removing these regions as sometimes suggested in the literature (Marquez-Luna *et al*. 2020; Lloyd-Jones *et al*. 2019) since these regions, especially the HLA region, contain many variants associated with many traits, and are therefore very useful for prediction. Other modifications, such as a better handling of numerical errors when working with exponentials (equation (6)), or a larger hyper-parameter search space, may also contribute to the improved robustness in LDpred2.

In LDpred2, we also expand the grid of hyper-parameters examined with now more values for *p* (21 instead of 7 by default in LDpred1) and for *h*^2^ (3 instead of 1). When testing the grid of hyper-parameters of *p* and *h*^2^, we also allow for testing an option to enable sparse models in LDpred2 (see below). Overall, we test a grid of 126 different values in LDpred2 instead of 7 in LDpred1. We also use a larger window size for computing correlations between variants (see above), yet LDpred2 is still as fast as LDpred1. The efficiency of LDpred2 is achieved through an efficient parallel implementation in C++. Efficient parallelization over the grid of hyper-parameters is possible because we use an on-disk sparse matrix format accessed using memory mapping. This special data format is available in R package bigsparser, which has been developed for this paper. As an example, it takes less than three hours in total to run LDpred2-inf, LDpred2-grid (126 hyper-parameter values parallelized over 8 cores) and LDpred2-auto (30 initial values for *p* parallelized over 8 cores) for a chromosome with 100,000 variants. It takes less than 36 hours to run all these LDpred2 models over 1.1 million HapMap3 variants at once (genome-wide). It takes 11 minutes to pre-compute the LD matrix for 10,000 individuals and 100,000 variants (using 16 cores).

LDpred2 also extends the original LDpred model in two ways. First, we provide a sparse option in LDpred2-grid which provides models that truly encourage sparsity, in contrast to LDpred1 which outputs very small non-zero effect sizes (Janssens and Joyner 2019). In practice, the sparse version of LDpred2 can drastically reduce the number of variants used in the PGS without impacting its predictive performance, as opposed to discarding the smallest effects after having run LDpred (as tested in Bolli *et al*. (2019)). The second extension is LDpred2-auto, which automatically estimates values for hyper-parameters *p* and *h*^2^, which therefore does not require any validation set to tune hyper-parameters from. LDpred2-auto is an attractive option in many applications, especially since we now also provide an LD reference that can be used directly. LDpred2-auto almost always performs equally well as LDpred2-grid in simulations as well as in real data applications.

However, LDpred2-auto requires that some quality control is performed on summary statistics (see Methods section “Quality control of summary statistics”). This quality control aims at ensuring that effects are transferable from the external GWAS summary statistics to the data where PGS are computed. Note that this quality control is also beneficial for other methods such as lassosum (data not shown). Even when this quality control is performed, LDpred2-auto can slightly underperform in some situations, e.g. for CAD and Asthma here (Figure 4). When looking more closely at the results for CAD, even though chains seemed to have converged (Figure S5), we can see that the heritability estimate from LDpred2-auto is off compared to the one from LD score regression (Table S5), and that its estimate of *p* (∼0.001) is off compared to the best performing *p* in LDpred2-grid (∼0.01 in figure S4). One possible explanation for this discrepancy is that the summary statistics we use for CAD come from a meta-analysis of 48 small studies, with some of them from non-European ancestries. Moreover, the summary statistics we use for Asthma come from a meta-analysis of 66 small multiancestry studies. This could break some of the assumptions used in LDpred2-auto and explain why LDpred2-auto underperforms for CAD and Asthma compared to LDpred2-grid. Future work is needed to study how to make best use of these multiancestry meta-analyses from many small studies in the context of genetic prediction. Future work is also needed to assess how relevant is the estimation of parameters from LDpred2-auto, and whether its estimates and its predictive performance could be improved by e.g. incorporating functional annotations.

In conclusion, we have shown that LDpred2 provides a better, faster and more robust implementation of the LDpred model. When compared to recently derived methods that showed higher predictive performance than LDpred1, LDpred2 provides the highest predictive performance for all the real traits tested here. For now, we recommend using the same HapMap3 variants used in PRS-CS and used here when running LDpred2. Indeed, HapMap3 variants have passed a number of quality controls, are generally well imputed and offer a good coverage of the whole genome. However, investigating alternatives in variant selection for LDpred2 and other PGS methods is a direction of future research for us. Would it be beneficial e.g. to use a set enriched for statistically significant variants?

## 4 Methods

### Simulation analyses

We use the UK Biobank imputed data for both real data analyses and simulations (Bycroft *et al*. 2018). We use dosage data for LDpred2, which we read from BGEN files using function `snp_readBGEN` from R package bigsnpr (Privé *et al*. 2018). For the other software that require a PLINK bed file as input, we use function `snp_writeBed` that round dosages to write genotype data in bed format. We restrict individuals to the ones used for computing the principal components (PCs) in the UK Biobank; these individuals are unrelated and have passed some quality control (Bycroft *et al*. 2018). To get a set of genetically homogeneous individuals, we compute a robust Mahalanobis distance based on the first 16 PCs and further restrict individuals to those within a log-distance of 5 (Privé *et al*. 2020). We restrict variants to the HapMap3 variants used in PRS-CS (Ge *et al*. 2019). This results in 362,320 individuals and 1,117,493 variants. We use 10,000 individuals as validation set for choosing optimal hyper-parameters and for computing correlations between variants (LD matrix ***R***). We use 300,000 other individuals for running logistic GWAS to create summary statistics. We use the remaining 52,320 individuals as test set for evaluating models.

We simulate binary phenotypes with a heritability of *h*^2^ = 0.4 using a Liability Threshold Model (LTM) with a prevalence of 15% (Falconer 1965). We vary the number of causal variants (300, 3000, 30,000, or 300,000) to match a range of genetic architectures from low to high polygenicity. Causal variants are chosen randomly anywhere on the genome. Liability scores are computed from a model with additive effects only: we compute the liability score of the *i*-th individual as 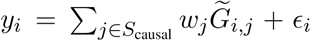, where *S*_causal_ is the set of causal variants, *w*_*j*_ are weights generated from a Gaussian distribution *N* (0, *h*^2^*/*|*S*_causal_|), *G*_*i,j*_ is the allele dosage of individual *i* for variant *j*, 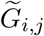 corresponds to its standardized version (zero mean and unit variance), and ϵ_*i*_ follows a Gaussian distribution *N* (0, 1 − *h*^2^). Both parts of the *y*_*i*_’s are scaled such that the variance of the genetic liability is exactly *h*^2^ and the variance of the total liability is exactly 1.

Such simulation of phenotypes based on real genotypes is implemented in function `snp_simuPheno` of R package bigsnpr. We also vary the sample size to compute GWAS summary statistics in the scenario with 3000 causal variants; in addition to a GWAS sample size of 300,000, we also use 10,000, 20,000, 50,000 and 120,000. We design two other simulation scenarios with 300 or 3000 causal variants randomly chosen in the HLA region (chromosome 6, 25.5-33.5 Mb). In these two scenarios, we use *h*^2^ = 0.3 instead of *h*^2^ = 0.4 because the total heritability is gathered in one chromosome only. Finally, we design a seventh simulation scenario as a mixture of previous scenarios; we simulate 300 causal variants in the HLA region explaining 20% of the variance in liability, and 10,000 causal variants anywhere on the genome explaining another 20% (*h*^2^ = 0.4 in total).

All simulation scenarios are summarized in table 1. Each simulation scenario is repeated 10 times and averages of the Area Under the ROC Curve (AUC) are reported. The 95% confidence interval (CI) from 10,000 non-parametric bootstrap replicates of the mean AUC of the 10 simulations for each scenario is also reported. In other words, we sample 10,000 bootstrap replicates of these 10 AUC values and compute their respective mean. We then report the mean of these 10,000 values, along with their quantile at 2.5% and at 97.5% to act as the 95% confidence interval (CI) for the mean AUC.

### Real data analyses

We use the same data as in the simulation analyses. We use the same 10,000 individuals as validation set, and use the remaining 352,320 individuals as test set. We use external published GWAS summary statistics listed in table 2. We defined phenotypes as in Privé *et al*. (2019). Some quality control is applied to summary statistics (see Method section “Quality control of summary statistics” below). For more details, please refer to our R code (Software and code availability section).

In real data applications, we first compare all four LDpred2 models to the two LDpred1 models. Then, we compare LDpred2-inf, LDpred2-grid and LDpred2-auto to several other methods: Clumping and Thresholding (C+T), Stacked C+T (SCT), lassosum(-auto), PRS-CS(-auto) and SBayesR (Privé *et al*. 2019; Mak *et al*. 2017; Ge *et al*. 2019; Lloyd-Jones *et al*. 2019). For C+T and SCT (Privé *et al*. 2019), we use the default large grid of hyper-parameters testing a threshold of clumping 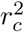 within {0.01, 0.05, 0.1, 0.2, 0.5, 0.8, 0.95}, a base size of clumping window within {50, 100, 200, 500} in Kb where the actual window size is then computed as the base size divided by 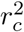, and a sequence of 50 thresholds on -log10(p-values) between 0.1 and the most significant p-value, equally spaced on a log scale. For lassosum (Mak *et al*. 2017), we use the default grid of hyper-parameters *s* and *λ*. We also alternatively choose the optimal hyper-parameters based on pseudo-validation (instead of choosing them based on performance in the validation set) and refer to this method as lassosum-auto. For PRS-CS (Ge *et al*. 2019), we use the grid {10^−6^, 10^−5^, 10^−4^, 10^−3^, 0.01, 0.1, 1} for the global scaling hyper-parameter Φ, and the default values for hyper-parameters a (1) and b (0.5). PRS-CS-auto automatically estimates Φ. Finally, for SBayesR (Lloyd-Jones *et al*. 2019), shrunk LD matrices are built using option ‘`--make-shrunk-ldm`’ and SBayesR is run with parameters ‘`--pi 0.95,0.02,0.02,0.01`’ and ‘`--gamma 0.0,0.01,0.1,1`’, and a chain of length 10,000 with 2000 burn-in iterations. Note that we use the 10,000 individuals of the validation set as LD reference for all methods, except for PRS-CS which provides its own LD reference based on 503 individuals from the 1000 Genomes data; Ge *et al*. (2019) argued that the sample size of the LD reference has little impact on the performance of PRS-CS.

We use the Area Under the ROC Curve (AUC) to compare methods. We sample 10,000 bootstrap replicates of the individuals in the test set and compute the AUC for each of these. We then report the mean of these 10,000 values, along with their quantile at 2.5% and at 97.5% to act as the 95% confidence interval (CI) for the AUC. This is implemented in function `AUCBoot` of R package bigstatsr.

### From marginal effects to joint effects

In this section, we explain how we can obtain joint effects from summary statistics (marginal effects) and a correlation matrix ***R***. Let us denote by ***S*** the diagonal matrix with standard deviations of the *m* variants, ***C***_***n***_ = ***I***_***n***_ − **11**^*T*^ */n* the centering matrix, ***G*** the genotype matrix of *n* individuals and *m* variants, and ***y*** the phenotype vector for *n* individuals.

When solving a joint model with all variants and an intercept *α*, the joint effects ***γ***_joint_ are obtained by solving

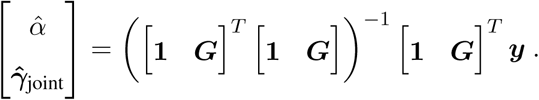

Using the Woodburry formula, we get

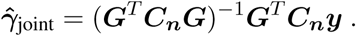

When fitting each variant separately in GWAS, the marginal effects (assuming no covariate) simplify to

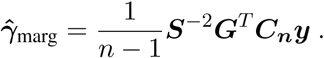

We further note that the correlation matrix of ***G*** is 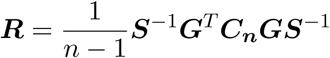. Then we get

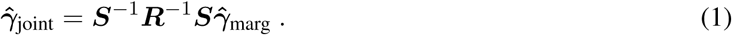

In practice, the correlation matrix ***R*** is usually not available but is computed from another dataset. Also note that ***γ*** are the effects on the allele scale while we denote by ***β*** = ***Sγ*** the effects of the scaled genotypes.

For the marginal effect 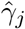 of variant *j*, let us denote by 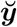 and 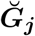 the vectors of phenotypes and genotypes for variant *j* residualized from *K* covariates, e.g. centering them. Then,

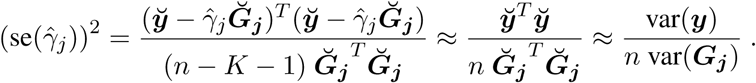

The first approximation is possible because 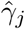 is expected to be small, while the second approximation assumes that the effects from covariates are small. Thus we can derive

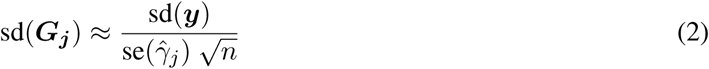

and then 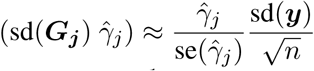. Let us go back to equation 1. As sd (***y***) is the same for all variants, it is cancelled out by ***S***^−1^ and ***S***, therefore we can assume that var(***y***) = 1. It justifies the use of the Z-scores 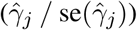 divided by 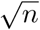 as input for LDpred (first line of algorithm 1). Then, the effect sizes that LDpred outputs need to be scaled back by multiplying by 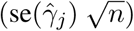 (last line of algorithm 1). In LDpred2, we allow for having different *n*_*j*_ for different variants. Note that LDpred1 and other similar methods scale the output dividing by the standard deviation of genotypes. This is correct when var(***y***) = 1 only.

### Quality control of summary statistics

For summary statistics of binary traits derived from a logistic regression, instead of equation (2), we have

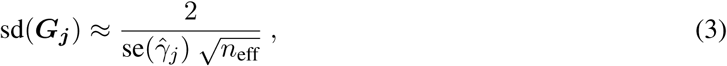

Where 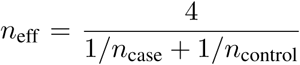. We recommend to verify this assumption and to perform some quality control. Indeed, in simulations, the approximation of equation (3) seems valid (Figure S6). However, in real data applications, where summary statistics come from a meta-analysis of many external datasets, this approximation can be invalidated (Figure S7). Let us denote by SD_ss_ the standard deviations derived from the summary statistics (right-hand side of equation (2) or (3)) and by SD_val_ the standard deviations of genotypes of individuals in the validation set (left-hand side). Note that, in order to compute SD_ss_ in the case of summary statistics from a linear regression, sd(***y***) from equation (2) can be estimated using e.g. the median value of 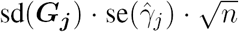 or by using the fact that the maximum value of SD_ss_ should be 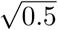. We recommend removing variants with SD_ss_ *<* 0.5 · SD_val_ or SD_ss_ > 0.1 + SD_val_ or SD_ss_ *<* 0.1 or SD_val_ *<* 0.05 (Figure S7).

### Overview of LDpred model

LDpred assumes the following model for effect sizes,

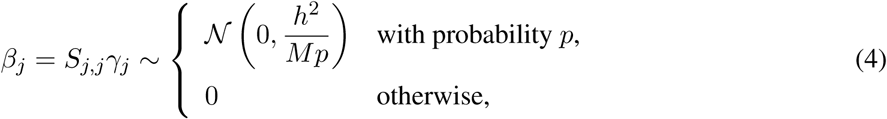

where *p* is the proportion of causal variants, *M* the number of variants and *h*^2^ the (SNP) heritability. Vilhjálmsson *et al*. (2015) estimate *h*^2^ using constrained LD score regression (intercept fixed to 1) and recommend testing a grid of hyper-parameter values for *p* (1, 0.3, 0.1, 0.03, 0.01, 0.003 and 0.001).

To estimate effect sizes *β*_*j*_, we use a Gibbs sampler as in Vilhjálmsson *et al*. (2015), which is described in algorithm 1. First, the residualized marginal effect for variant *j* is computed as

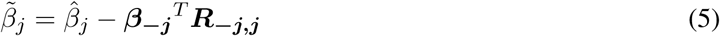

where ***R***_***−j*,*j***_ is the *j*-th column without the *j*-th row of the correlation matrix, 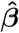 is the vector of marginal effect sizes, ***β*** is the vector of current effect sizes in the Gibbs sampler, and ***β***_***−j***_ is ***β*** without the *j*-th element. Then, the probability that variant *j* is causal is computed as

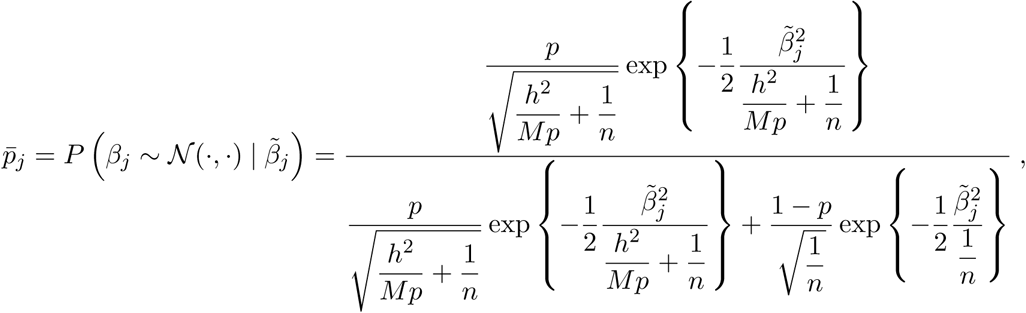

which we rewrite as

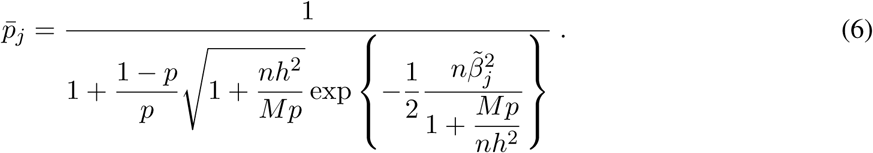

Computing 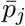 using the second expression is important to avoid numerical issues when 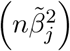 is large.

Then, *β*_*j*_ is sampled according to

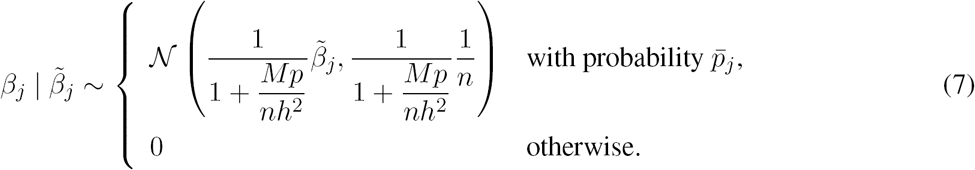

Therefore, the posterior mean of 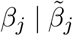 is given by

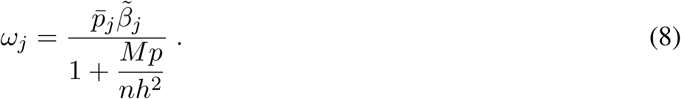

#### Algorithm 1

LDpred, with hyper-parameters *p* and *h*^2^, LD matrix ***R*** and summary statistics 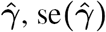 and ***n***

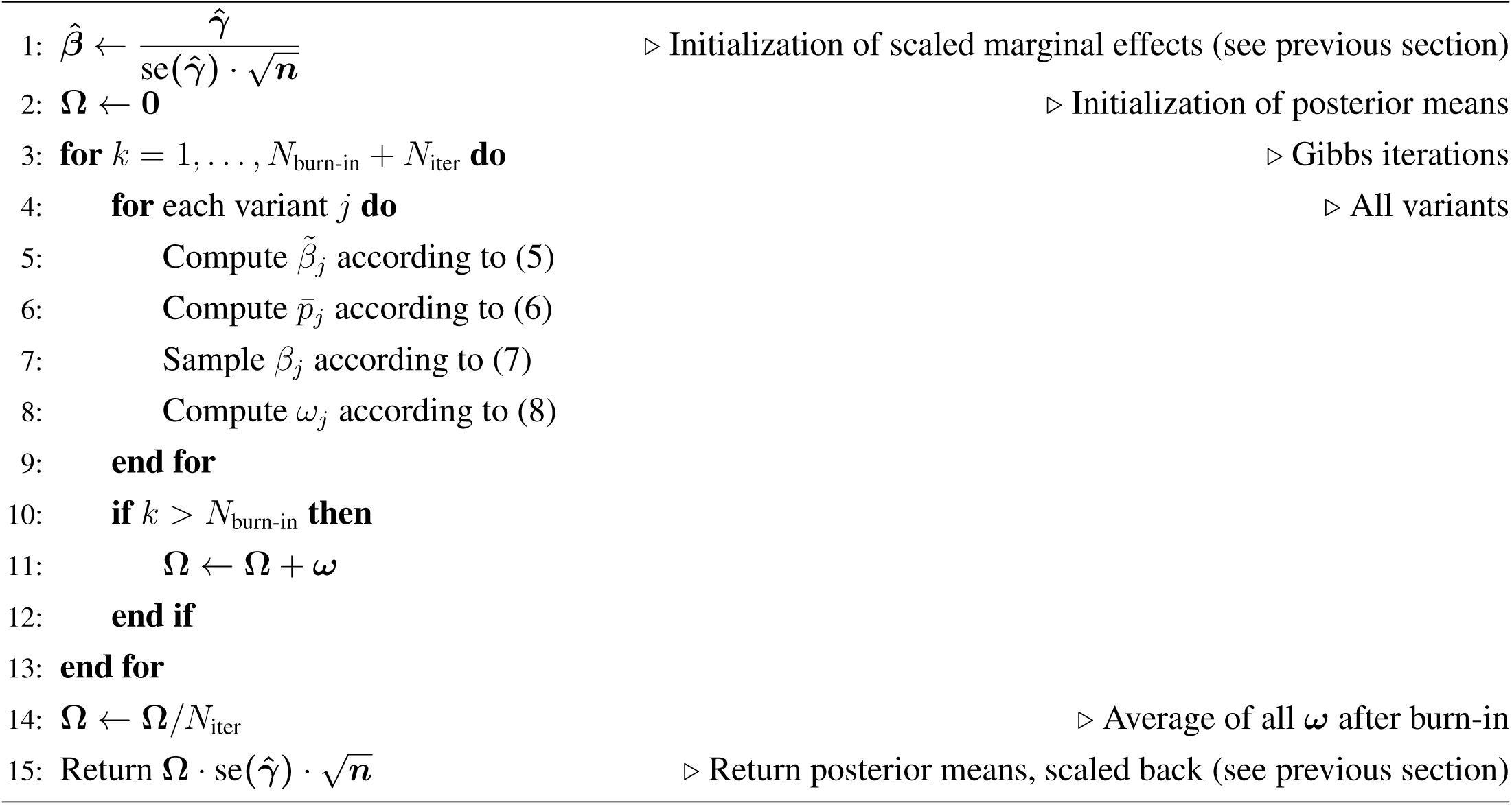

### New LDpred2 models

LDpred2 comes with two extensions of the LDpred model.

The first extension consists in estimating *p* and *h*^2^ within the model, as opposed to testing several values of *p* and estimating *h*^2^ using constrained LD score regression (Bulik-Sullivan *et al*. 2015). This makes LDpred2-auto a method free of hyper-parameters which can therefore be applied directly to data without the need of a validation dataset to choose best-performing hyper-parameters. To estimate *p* in the Gibbs sampler, we count the number of non-zero variants (i.e. *M*_*c*_ = ∑_*j*_ (*β*_*j*_ 0) in equation 7). We can assume that *M*_*c*_ ∼ Binom(*M, p*), so if we place a prior *p* ∼ Beta(1, 1) ≡ 𝒰(0, 1), we can sample *p* from the posterior *p* ∼ Beta(1 + *M*_*c*_, 1 + *M* − *M*_*c*_). Due to complexity reasons, we could not derive a Bayesian estimator of *h*^2^. Instead, we estimate *h*^2^ = ***β***^*T*^ ***Rβ***, where ***R*** is the correlation matrix. These parameters *p* and *h*^2^ are updated after the inner loop in algorithm 1, then these new values are used in the next iteration of the outer loop.

The second extension, which can be enabled using a third hyper-parameter in LDpred2-grid, aims at providing sparse effect size estimates, i.e. some resulting effects are exactly 0. When the sparse solution is sought and when 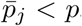, we set *β*_*j*_ and *ω*_*j*_ to 0 (lines 6-8 of algorithm 1). We also provide a sparse option for LDpred2-auto by running LDpred2-grid with one set of parameters only: with the sparsity enabled and using the estimates of *p* and *h*^2^ from LDpred2-auto.

When running LDpred2-grid, we test a grid of hyper-parameters with *p* from a sequence of 21 values from 10^−5^ to 1 on a log-scale; *h*^2^ within {0.7, 1, 1.4} · 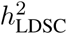, where 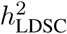 is the heritability estimate from the constrained LD score regression Bulik-Sullivan *et al*. (2015); and whether sparsity is enabled or not. In total, this grid is of size 21 × 3 × 2 = 126. When running LDpred2-auto, we run it 30 times with 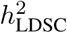 as initial value for *h*^2^ and a sequence of 30 values from 10^−4^ to 0.9 equally spaced on a log scale as initial values for *p*. Running many Gibbs chains aims at checking whether models did not diverge (and hopefully converged). As a criterion for non-divergence, we compute the standard deviations of the resulting predictors from the 30 models, keep only those within 3 median absolute deviations from their median, and average the remaining vectors of effects as final effect sizes for the “auto” version.

### New strategy for local correlation

There is a window size parameter that needs to be set in LDpred; for a given variant, correlations with other variants outside of this window are assumed to be 0. The recommended value for this window (in number of variants) has been to use the total number of variants divided by 3000, which corresponds to a window radius of around 2 Mb (Vilhjálmsson *et al*. 2015). We have come to the conclusion that this window size is not large enough. Indeed, the human leukocyte antigen (HLA) region of chromosome 6 is 8 Mb long (Price *et al*. 2008). Using a window of 8Mb would be computationally and memory inefficient. Instead, we propose to use genetic distances. Genome-wide, 1 Mb corresponds on average to 1 cM. Yet, the HLA region is only 3 cM long (vs. 8 Mb long). Therefore, genetic distances enable to capture the same LD using a globally smaller window. We provide function `snp_asGeneticPos` in package bignspr to easily interpolate physical positions (in bp) to genetic positions (in cM). We recommend to use genetic positions and to use a size parameter of 3 cM when computing the correlation between variants for LDpred2. Note that, in the code, we use ‘`size = 3 / 1000`’ since parameter `size` is internally multiplied by 1000 in the bigsnpr functions.

### Running LDpred2 per chromosome or genome-wide

In this paper, we investigate if it would be beneficial to run LDpred2 per chromosome, instead of genome-wide. Indeed, running LDpred2 genome-wide has two drawbacks. First, it can be memory and computationally demanding to do so. For around one million (1M) variants, storing the 1M×1M sparse correlation matrix takes more than 32 GB of memory. Doubling to 2M variants would require 128 GB of RAM to store the matrix. Second, it may be beneficial to assume that the architecture of traits is different for different chromosomes. For example, chromosome 6 clearly encompasses a larger proportion of the heritability of autoimmune diseases compared to other chromosomes (Shi *et al*. 2016). Assuming the same model for genetic effects genome-wide could result in model misspecification, which would lead to suboptimal predictive performance. To choose the best LDpred2 model, one can choose the best model according to their preferred criterion (e.g. max AUC). Here, we use the Z-Score from the regression of the phenotype by the PRS since we have found it more robust than using the AUC when running LDpred2 per chromosome.

### Providing an LD reference

We provide an LD reference for European ancestry to be used by researchers who cannot compute their own. We use the 362,320 UKBB individuals as used here, with some further quality control based on allele frequencies. Indeed, some large allele frequency mismatches have been reported recently in the UKBB when comparing to other datasets, which are apparently due to mismappings (Kunert-Graf *et al*. 2020). Note that most of these errors should be captured by the quality control we propose in this paper. Using 503 European individuals from the 1000 Genomes (1000G) data, we remove variants with allele frequency differences between UKBB and 1000G at *p <* 10^−5^ (Figure S8). We also remove variants with minor allele frequencies less than 1% in the 1000G or less than 0.5% in the UKBB. 1,054,330 variants remain, which we use to compute the LD reference provided. We also provide an example R script on how to use this LD reference provided.

## Software and code availability

The newest version of R package bigsnpr can be installed from GitHub (see https://github.com/privefl/bigsnpr). A tutorial on the steps to run LDpred2 using some small example data is available at https://privefl.github.io/bigsnpr/articles/LDpred2.html. The European LD reference we provide here is available at https://doi.org/10.6084/m9.figshare.13034123. All code used for this paper is available at https://github.com/privefl/paper-ldpred2/tree/master/code. We have extensively used R packages bigstatsr and bigsnpr (Privé *et al*. 2018) for analyzing large genetic data, packages from the future framework (Bengtsson 2020) for easy scheduling and parallelization of analyses on the HPC cluster, and packages from the tidyverse suite (Wickham *et al*. 2019) for shaping and visualizing results.

## Acknowledgements

Authors would like to thank the four reviewers of this paper for their relevant comments and valuable suggestions, Naomi Wray and Alkes Price for pointing to issues due to long-range LD regions in LDpred1, Yixuan Qiu for pointing to matrix-free solvers, Doug Speed and others for early testing of the software and for providing useful feedback, GenomeDK and Aarhus University for providing computational resources and support that contributed to these research results. This research has been conducted using the UK Biobank Resource under Application Number 41181.

