## Supplementary Tables and Figures for "LDpred2: better, faster, stronger"

### Simulations

| Method | all_40_300 | all_40_3000 | all_40_30000 | all_40_300000 | both_40 | HLA_30_300 | HLA_30_3000 |
|---|---|---|---|---|---|---|---|
| LDpred1-inf | 68.9 [68.4-69.3] | 69.3 [68.9-69.6] | 69.5 [69.2-69.7] | 69.5 [69.3-69.6] | 62.9 [61.6-64.3] | 54.4 [52.9-55.9] | 54.5 [53.7-55.5] |
| LDpred2-inf-perchr | 70.3 [69.9-70.6] | 70.0 [69.7-70.2] | 69.9 [69.7-70.1] | 69.9 [69.7-70.0] | 74.2 [73.8-74.5] | 75.6 [74.8-76.3] | 75.7 [75.4-75.9] |
| LDpred1-grid | 73.6 [72.0-75.4] | 70.4 [69.8-70.9] | 69.1 [68.8-69.3] | 69.2 [69.0-69.4] | 61.7 [60.4-63.1] | 54.7 [53.9-55.5] | 54.5 [53.8-55.1] |
| LDpred2-grid-nosp-perchr | 81.7 [81.6-81.9] | 78.9 [78.7-79.0] | 70.6 [70.4-70.8] | 69.3 [69.0-69.6] | 72.8 [67.3-75.7] | 76.4 [75.7-77.0] | 76.4 [76.1-76.6] |
| LDpred2-grid-sp-perchr | 81.7 [81.6-81.9] | 78.8 [78.7-78.9] | 70.5 [70.3-70.7] | 69.4 [69.1-69.6] | 72.8 [67.3-75.7] | 76.5 [75.8-77.1] | 76.5 [76.3-76.7] |
| LDpred2-auto-perchr | 82.0 [81.8-82.2] | 79.3 [79.2-79.4] | 70.8 [70.6-71.0] | 69.8 [69.6-70.0] | 73.7 [73.3-74.2] | 73.1 [72.0-74.1] | 72.9 [71.6-74.0] |

Table S1: Values from figure 1.

| Method | 10000 | 20000 | 50000 | 120000 | 300000 |
|---|---|---|---|---|---|
| LDpred1-grid | 56.1 [55.8-56.4] | 59.3 [58.9-59.8] | 66.3 [65.2-67.1] | 70.7 [69.2-71.7] | 70.4 [69.8-70.9] |
| LDpred1-inf | 55.7 [55.4-56.0] | 57.7 [57.4-58.0] | 61.2 [60.9-61.5] | 65.0 [64.7-65.3] | 69.3 [69.0-69.7] |
| LDpred2-auto-gwide | 55.9 [55.5-56.2] | 59.3 [58.9-59.8] | 67.5 [67.3-67.8] | 74.7 [74.6-74.9] | 79.3 [79.2-79.4] |
| LDpred2-auto-perchr | 54.1 [53.7-54.5] | 57.3 [56.8-57.8] | 66.6 [66.3-66.9] | 74.7 [74.5-74.8] | 79.3 [79.2-79.4] |
| LDpred2-grid-nosp-gwide | 55.7 [55.2-56.0] | 59.6 [59.3-60.0] | 67.4 [67.2-67.7] | 74.6 [74.5-74.8] | 79.1 [79.0-79.3] |
| LDpred2-grid-nosp-perchr | 54.4 [54.0-54.8] | 58.6 [58.1-59.1] | 66.8 [66.5-67.1] | 74.2 [73.9-74.5] | 78.9 [78.7-79.0] |
| LDpred2-grid-sp-gwide | 56.1 [55.7-56.5] | 59.6 [59.3-60.0] | 67.4 [67.2-67.7] | 74.6 [74.4-74.8] | 79.2 [79.1-79.3] |
| LDpred2-grid-sp-perchr | 54.5 [54.2-54.9] | 58.4 [57.9-58.9] | 66.8 [66.6-67.1] | 74.2 [73.9-74.4] | 78.8 [78.7-78.9] |
| LDpred2-inf-gwide | 55.5 [55.2-55.8] | 57.7 [57.5-58.0] | 61.4 [61.1-61.6] | 65.4 [65.1-65.6] | 69.9 [69.7-70.1] |
| LDpred2-inf-perchr | 54.6 [54.3-55.0] | 57.2 [56.8-57.5] | 61.3 [61.0-61.5] | 65.4 [65.2-65.6] | 70.0 [69.7-70.2] |

Table S2: Values from figures 2 and S2.

### Real data

| Method | Asthma | BRCA | CAD | MDD | PRCA | RA | T1D | T2D |
|---|---|---|---|---|---|---|---|---|
| LDpred2-grid-nosp-perchr | 58.8 [58.5-59.1] | 64.6 [64.1-65.1] | 61.5 [61.0-62.0] | 57.9 [57.5-58.2] | 68.2 [67.6-68.7] | 59.9 [59.2-60.6] | 77.0 [75.2-78.8] | 63.3 [62.9-63.7] |
| LDpred2-grid-sp-perchr | 58.5 [58.2-58.8] | 64.6 [64.1-65.0] | 61.2 [60.8-61.7] | 58.2 [57.8-58.6] | 67.8 [67.3-68.4] | 60.0 [59.3-60.7] | 75.9 [74.0-77.7] | 63.2 [62.7-63.6] |
| LDpred2-auto-perchr | 58.2 [57.9-58.5] | 65.6 [65.2-66.1] | 62.1 [61.6-62.6] | 58.9 [58.5-59.2] | 70.2 [69.6-70.7] | 59.7 [58.9-60.4] | 76.6 [74.7-78.5] | 63.9 [63.5-64.3] |
| LDpred2-inf-perchr | 57.1 [56.8-57.4] | 61.9 [61.4-62.4] | 60.8 [60.4-61.3] | 58.9 [58.5-59.2] | 65.0 [64.4-65.6] | 59.6 [58.9-60.3] | 74.5 [72.6-76.3] | 61.5 [61.1-61.9] |
| LDpred2-inf-gwide | 57.2 [56.9-57.4] | 61.6 [61.2-62.1] | 61.6 [61.1-62.1] | 59.0 [58.6-59.3] | 64.4 [63.8-65.0] | 59.3 [58.6-60.0] | 71.0 [69.1-72.8] | 61.6 [61.1-62.0] |
| LDpred2-grid-nosp-gwide | 59.3 [59.0-59.6] | 65.5 [65.0-66.0] | 63.6 [63.2-64.1] | 59.1 [58.7-59.4] | 70.2 [69.6-70.7] | 60.3 [59.6-61.0] | 78.4 [76.7-80.2] | 64.3 [63.8-64.7] |
| LDpred2-grid-sp-gwide | 59.4 [59.1-59.7] | 65.7 [65.3-66.2] | 63.5 [63.1-64.0] | 59.0 [58.7-59.4] | 70.2 [69.6-70.7] | 60.5 [59.8-61.3] | 78.3 [76.5-80.0] | 64.0 [63.6-64.5] |
| LDpred2-auto-gwide | 58.4 [58.2-58.7] | 65.6 [65.2-66.1] | 61.8 [61.3-62.3] | 59.0 [58.6-59.4] | 70.1 [69.5-70.7] | 59.7 [59.0-60.4] | 77.7 [75.9-79.4] | 63.8 [63.3-64.2] |
| LDpred1-grid | 58.6 [58.3-58.8] | 58.9 [58.4-59.4] | 60.3 [59.9-60.8] | 56.4 [56.1-56.8] | 63.5 [62.9-64.1] | 59.1 [58.4-59.8] | 57.4 [55.5-59.2] | 50.3 [49.8-50.7] |
| LDpred1-inf | 56.9 [56.6-57.1] | 58.8 [58.3-59.3] | 59.3 [58.9-59.8] | 56.5 [56.1-56.8] | 61.6 [61.0-62.3] | 59.5 [58.8-60.2] | 57.7 [55.8-59.5] | 51.8 [51.4-52.2] |

Table S3: Values from figures 3 and S3.

| Method | Asthma | BRCA | CAD | MDD | PRCA | RA | T1D | T2D |
|---|---|---|---|---|---|---|---|---|
| LDpred2-inf-gwide | 57.2 [56.9-57.4] | 61.6 [61.1-62.1] | 61.6 [61.1-62.0] | 59.0 [58.6-59.3] | 64.4 [63.8-65.0] | 59.3 [58.6-60.0] | 71.0 [69.1-72.9] | 61.6 [61.1-62.0] |
| LDpred2-grid-nosp-gwide | 59.3 [59.0-59.6] | 65.5 [65.0-66.0] | 63.6 [63.2-64.1] | 59.1 [58.7-59.4] | 70.2 [69.6-70.8] | 60.3 [59.6-61.0] | 78.4 [76.7-80.1] | 64.3 [63.9-64.7] |
| LDpred2-grid-sp-gwide | 59.4 [59.1-59.7] | 65.7 [65.3-66.2] | 63.5 [63.1-64.0] | 59.0 [58.7-59.4] | 70.2 [69.6-70.7] | 60.5 [59.8-61.3] | 78.2 [76.5-80.0] | 64.0 [63.6-64.5] |
| LDpred2-auto-gwide | 58.4 [58.2-58.7] | 65.6 [65.2-66.1] | 61.8 [61.3-62.3] | 59.0 [58.6-59.4] | 70.1 [69.5-70.7] | 59.7 [59.0-60.5] | 77.7 [75.9-79.4] | 63.8 [63.3-64.2] |
| SCT | 58.3 [58.0-58.6] | 62.3 [61.8-62.8] | 60.6 [60.1-61.1] | 57.4 [57.1-57.8] | 66.5 [65.9-67.1] | 57.4 [56.7-58.1] | 72.4 [70.5-74.2] | 62.5 [62.0-62.9] |
| C+T | 56.7 [56.4-57.0] | 62.9 [62.5-63.4] | 61.6 [61.1-62.1] | 58.5 [58.1-58.9] | 67.3 [66.7-67.9] | 59.1 [58.4-59.8] | 74.4 [72.6-76.2] | 59.9 [59.4-60.3] |
| lassosum | 57.6 [57.3-57.8] | 65.2 [64.7-65.6] | 62.5 [62.1-63.0] | 59.1 [58.8-59.5] | 69.3 [68.7-69.9] | 59.3 [58.5-59.9] | 74.3 [72.5-76.1] | 62.7 [62.3-63.2] |
| lassosum-auto | 57.6 [57.3-57.8] | 65.1 [64.7-65.6] | 61.4 [61.0-61.9] | 54.9 [54.6-55.3] | 69.3 [68.7-69.8] | 58.2 [57.5-58.9] | 75.8 [74.1-77.5] | 62.4 [62.0-62.8] |
| PRS-CS | 57.1 [56.8-57.4] | 63.3 [62.8-63.7] | 61.8 [61.4-62.3] | 57.5 [57.2-57.9] | 67.5 [66.9-68.1] | 59.2 [58.5-59.9] | 74.2 [72.4-76.0] | 62.2 [61.7-62.6] |
| PRS-CS-auto | 56.6 [56.3-56.8] | 63.4 [62.9-63.8] | 60.9 [60.4-61.3] | 53.9 [53.5-54.3] | 67.2 [66.6-67.7] | 58.6 [57.9-59.3] | 73.9 [72.1-75.7] | 62.4 [62.0-62.9] |
| SBayesR | 57.6 [57.4-57.9] | 65.7 [65.2-66.2] | 62.2 [61.7-62.6] | 58.8 [58.5-59.2] | 69.6 [69.1-70.2] | 56.2 [55.5-56.9] | 58.1 [56.1-60.2] | 64.1 [63.7-64.5] |

Table S4: Values from figures 4 and S1.

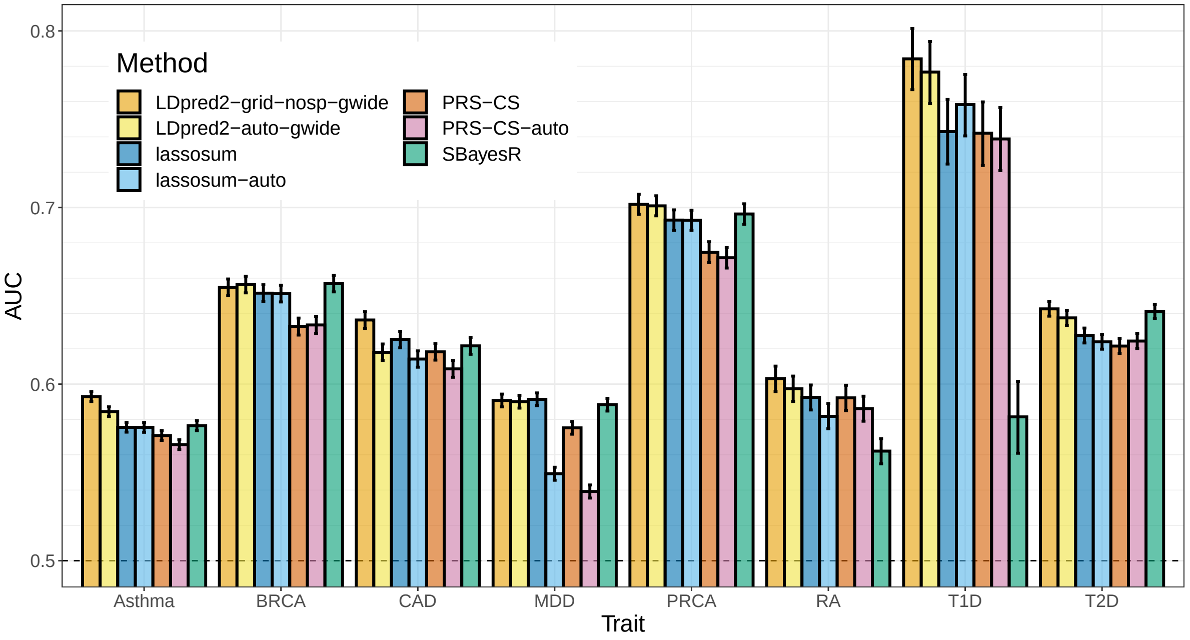


Figure S1: Both "grid" and "auto" variants are compared in real data applications for LDpred2, lassosum, PRS-CS and SBayesR. Note that SBayesR is such an "auto" model, but does not have any "grid" counterpart. Bars present AUC values on the test set of UKBB (mean and 95% CI from 10,000 bootstrap samples). See corresponding values in table S4.

### Running LDpred2 per chromosome or genome-wide?

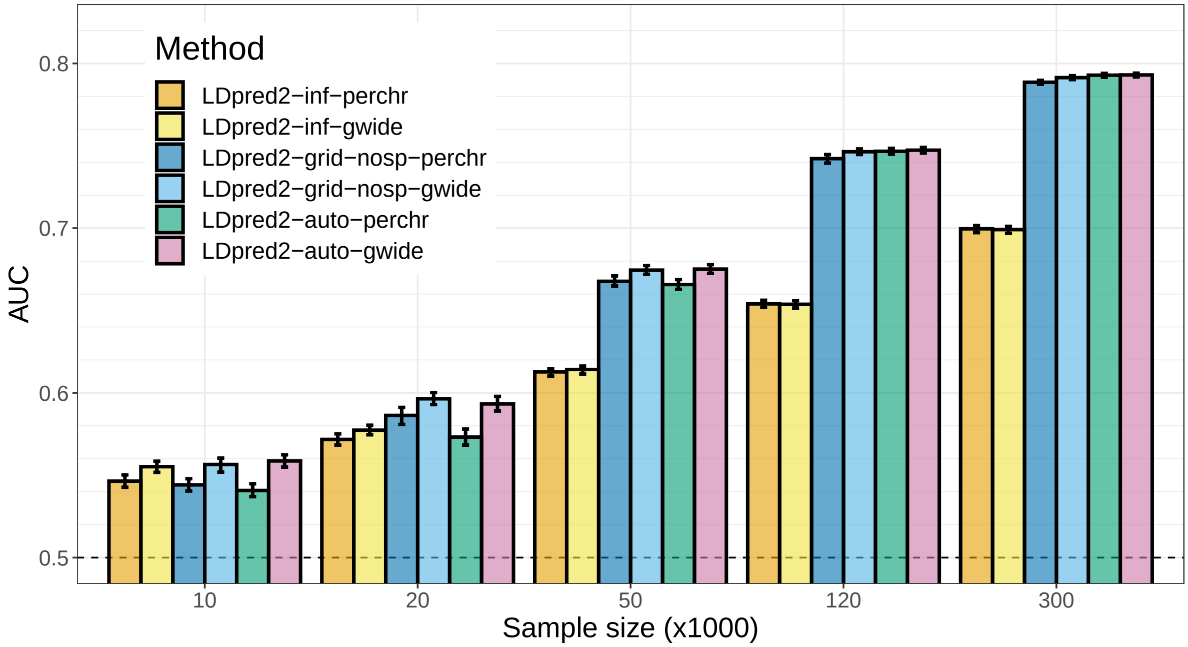


Figure S2: LDpred2 models, run either per chromosome or genome-wide, are compared when varying GWAS sample size in scenario “all_40_3000”. Bars present the mean and 95% CI of 10,000 non-parametric bootstrap replicates of the mean AUC of 10 simulations for each scenario. See corresponding values in table S2.

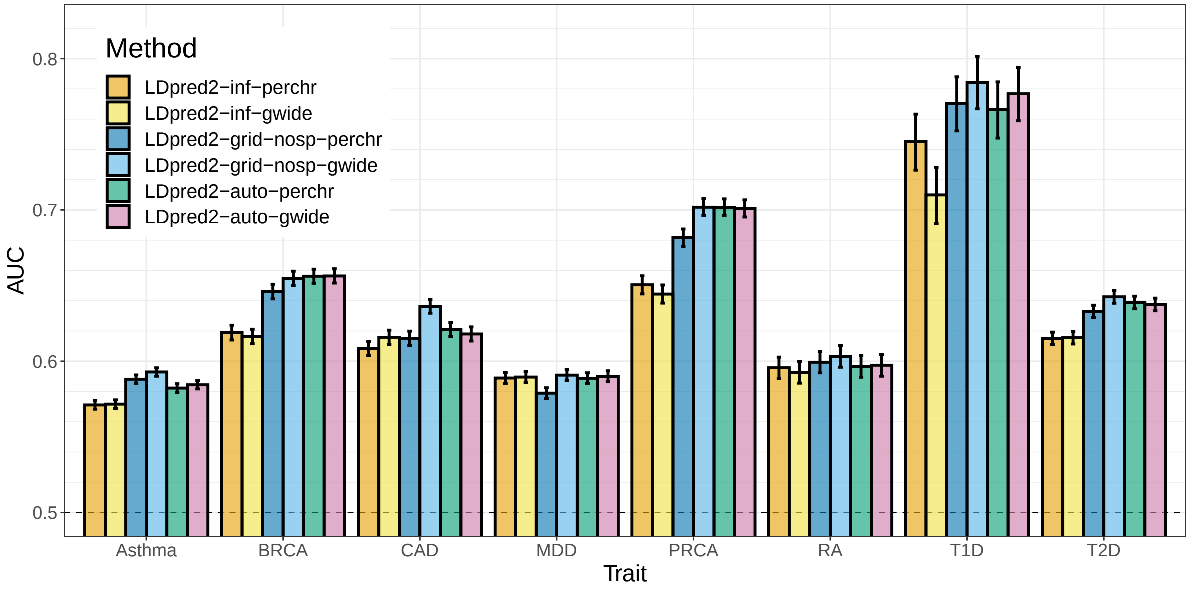


Figure S3: LDpred2 models, run either per chromosome or genome-wide, are compared in real data applications. Bars present AUC values on the test set of UKBB (mean and 95% CI from 10,000 bootstrap samples). See corresponding values in table S3.

### Estimation of parameters in LDpred2

| Trait | h2_ldsc_obs | sparsity_grid | p_auto | h2_auto_obs | prev_pop | h2_auto_liab |
|---|---|---|---|---|---|---|
| Asthma | 0.104 | 0.544 | 0.000738 | 0.0534 | 0.15 | 0.064 |
| BRCA | 0.127 | 0.414 | 0.00474 | 0.141 | 0.0758 | 0.136 |
| CAD | 0.0819 | 0.441 | 0.000978 | 0.0428 | 0.0619 | 0.0387 |
| MDD | 0.0923 | 0.377 | 0.0675 | 0.0877 | 0.0893 | 0.089 |
| PRCA | 0.188 | 0.516 | 0.00181 | 0.189 | 0.0546 | 0.164 |
| RA | 0.416 | 0.193 | 0.0774 | 0.562 | 0.0286 | 0.406 |
| T1D | 0.955 | 0.498 | 0.00482 | 1.4 | 0.00244 | 0.575 |
| T2D | 0.148 | 0.498 | 0.0031 | 0.131 | 0.0512 | 0.112 |

Table S5: Parameters obtained when running LDpred2 genome-wide for all 8 real traits analyzed here. h2_ldsc_obs: SNP heritability estimate (on the observed scale) from LD score regression (with intercept); sparsity_grid: sparsity of resulting effects from LDpred2-grid-sp; p_auto: estimate from LDpred2-auto of the proportion of causal variants; h2_auto_obs: SNP heritability estimate from LDpred2-auto; prev_pop: assumed population prevalence based on prevalence in the UK Biobank; h2_auto_liab: transformation of h2_auto_obs on the liability scale when assuming population prevalence prev_pop.

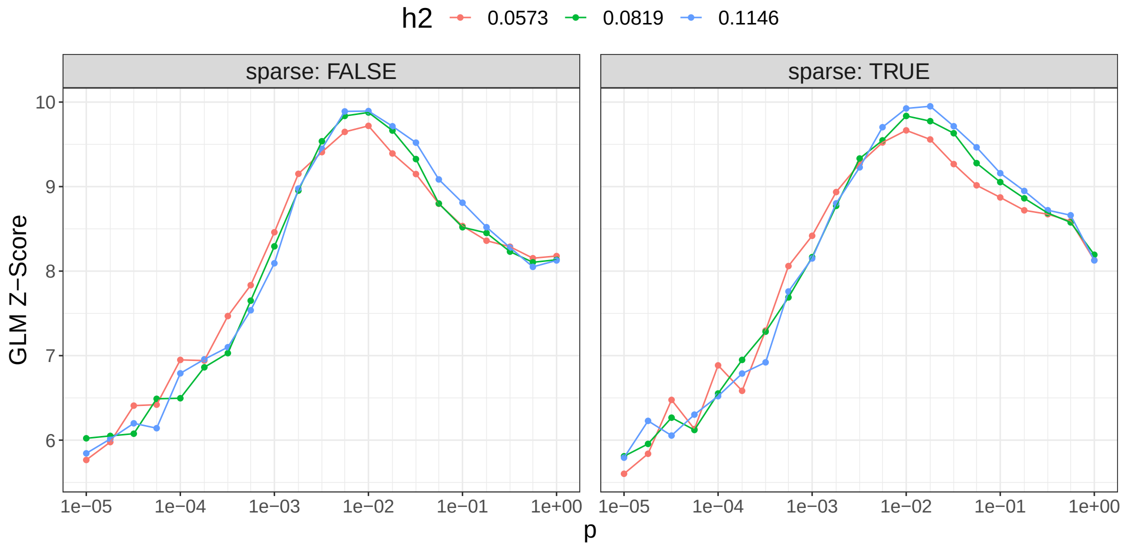


Figure S4: Results for all 126 parameter combinations of LDpred2-grid run genome-wide when applied to summary statistics for CAD.

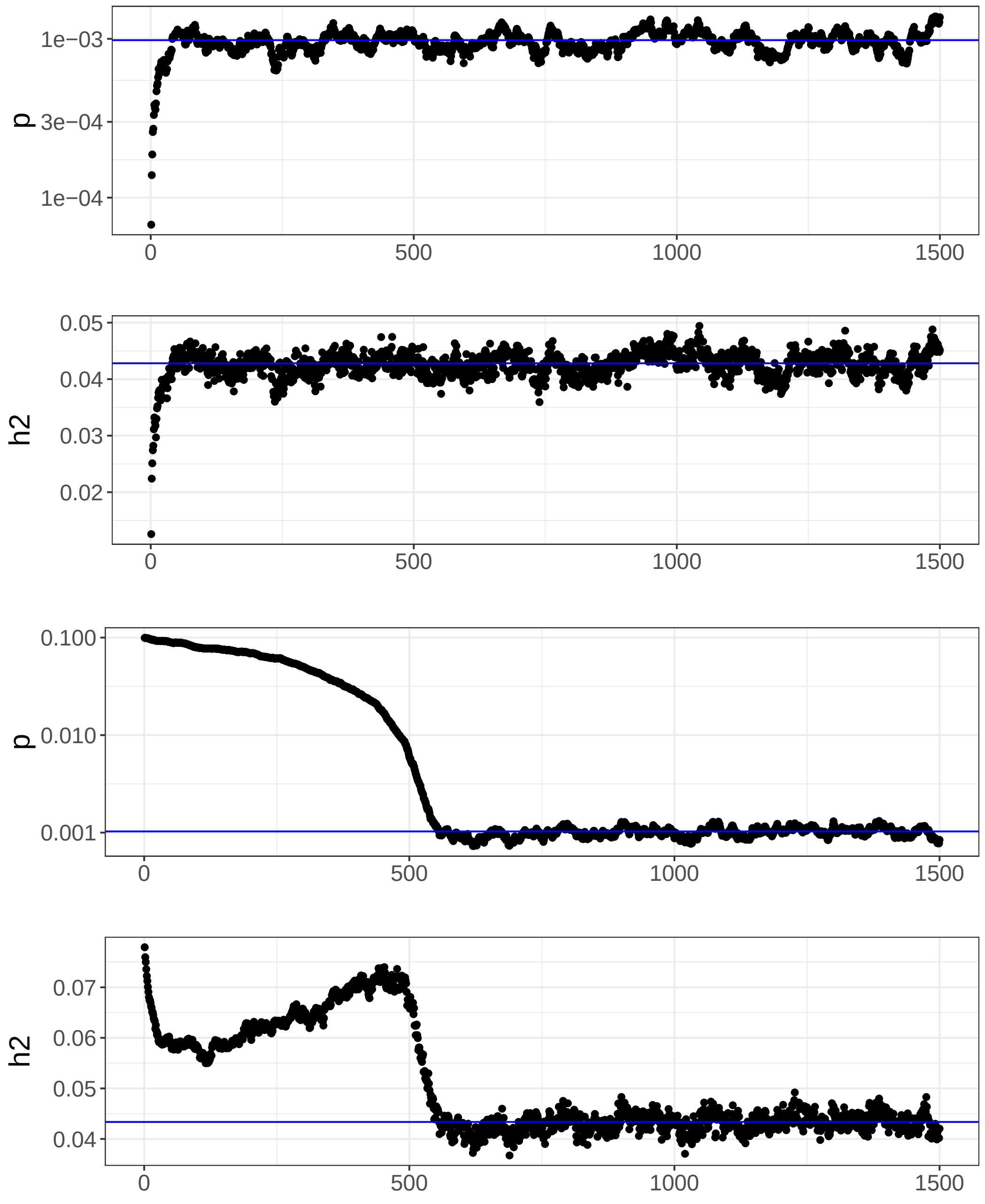


Figure S5: Paths of LDpred2-auto-gwide parameters, the SNP heritability $h^2$ and the proportion of causal variants $p$, for CAD. We show these for two different starting values for $p$, with an initial $p$ of 0.0001 at the top and initial $p$ of 0.1 at the bottom.

### Quality control of summary statistics

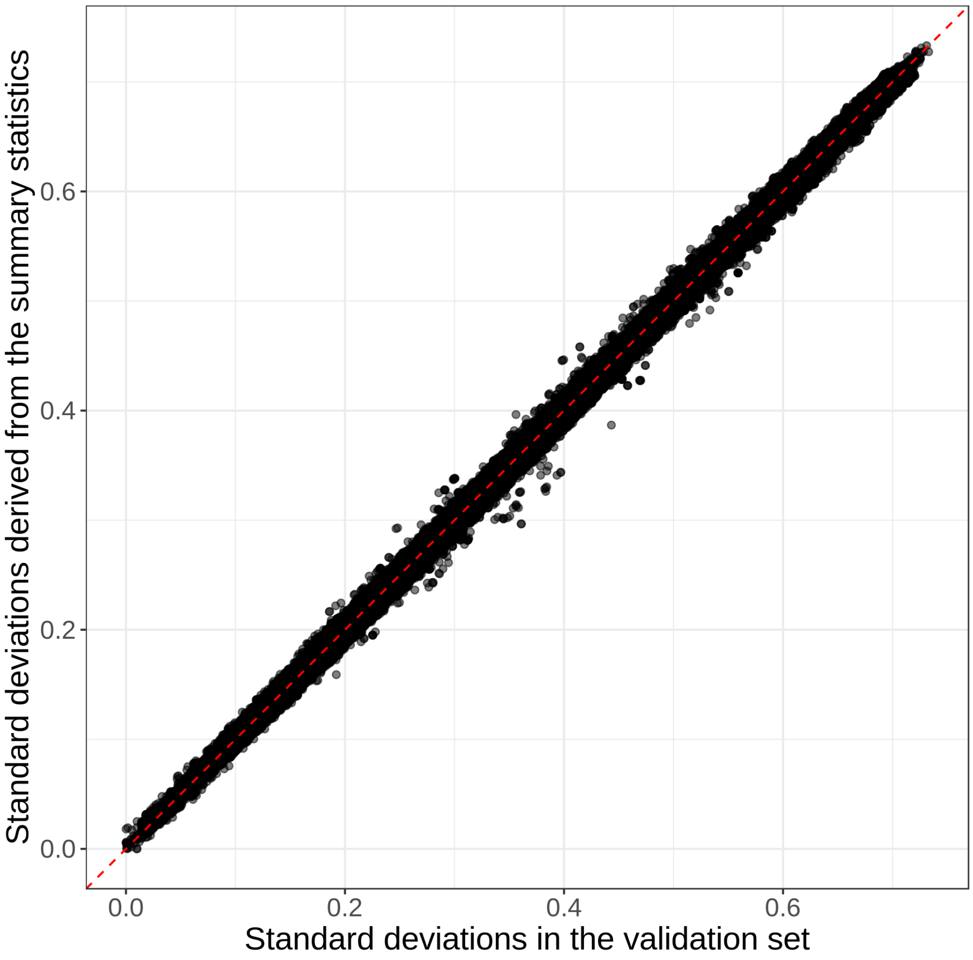


Figure S6: In simulations, standard deviations derived from summary statistics based on equation (3) versus the standard deviations of genotypes of individuals in the validation set.

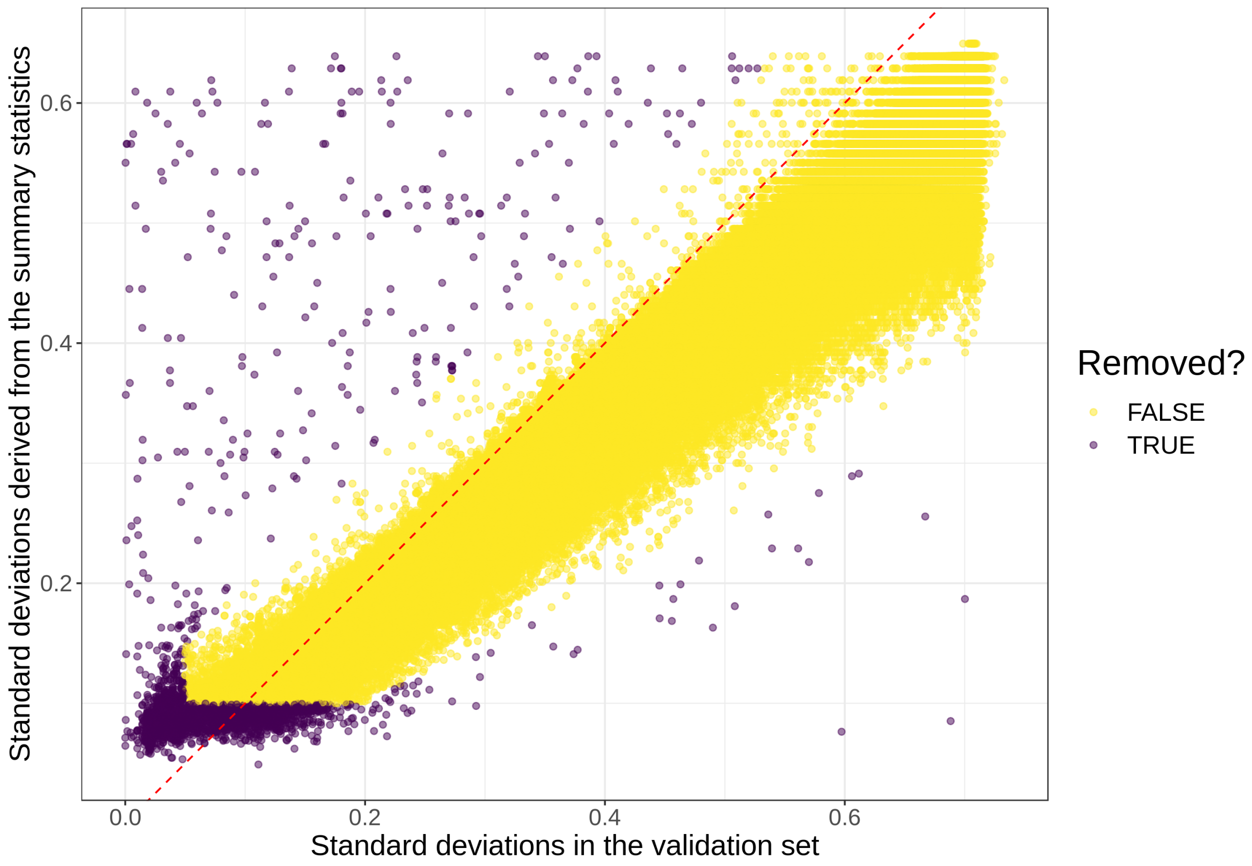


Figure S7: Standard deviations derived from summary statistics of breast cancer based on equation (3) versus the standard deviations of genotypes of individuals in the validation set. Coloring shows the quality control applied in this paper.

### Providing an LD reference

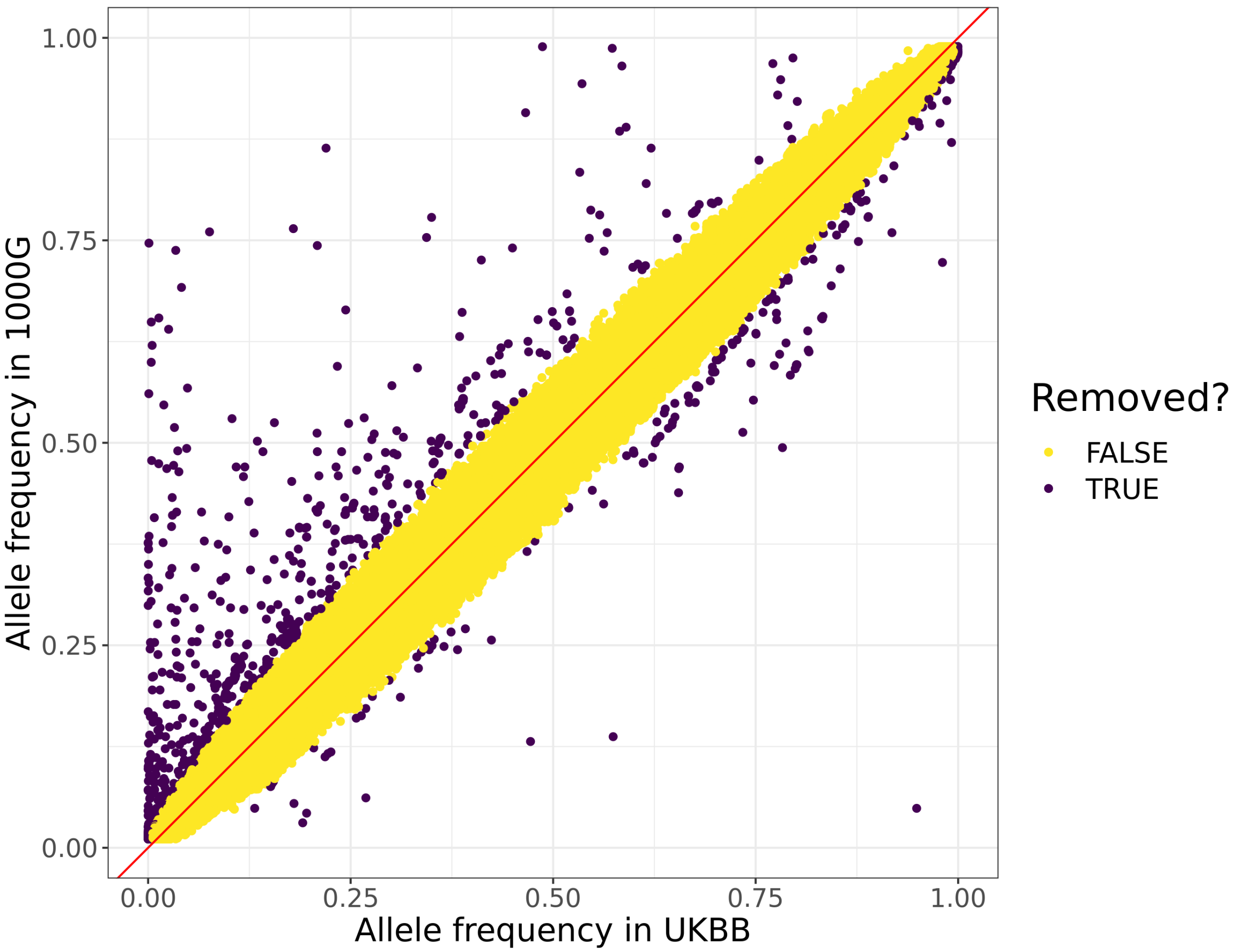


Figure S8: Allele frequencies for HapMap3 variants in the 1000 Genomes and in UK Biobank. Variants are removed when the difference between these two frequencies is very significant ($p < 10^{-5}$). Variants kept are used to provide an LD reference based on the UK Biobank data.
